# WHO child growth standards for Pygmies: one size fits all?

**DOI:** 10.1101/591172

**Authors:** Stephan M. Funk, Belén Palomo Guerra, Amy Ickowitz, Nicias Afoumpam Poni, Mohamadou Aminou Abdou, Yaya Hadam Sibama, René Penda, Guillermo Ros Brull, Martin Abossolo, Eva Ávila Martín, Robert Okale, Blaise Ango Ze, Ananda Moreno Carrión, Cristina García Sebastián, Cristina Ruiz de Loizaga García, Francisco López-Romero Salazar, Hissein Amazia, Idoia Álvarez Reyes, Rafaela Sánchez Expósito, John E. Fa

**Author notes:** joint first authors (Stephan M. Funk, Belén Palomo Guerra).

## Abstract

**Background:** African Pygmies exhibit a unique, genetically determined child growth dynamics and adult stature but the impact on assessing undernutrition remains unknown. Baka Pygmy health is highly compromised compared to sympatric populations. Evaluating child undernutrition is an important step to address this health quandry. We estimate stunting and wasting in Cameroon’s Baka children and investigate the applicability of the standards for Pygmy people.

**Methods:** Anthropometric and health data from 685 2-to12 year old children were collected at 25 health centres in southern Cameroon. Growth was analysed using both, WHO Child Growth Standards and the population itself as reference to define frequencies of stunting, wasting and obesity.

**Findings:** Baka children revealed with 68.4% the highest recorded level globally of stunting relative to the WHO child growth standard in 2-to-4 year olds. Wasting was at 8.2% in the upper third range in Sub-Saharan Africa. Obesity was with 6.5% similar to wasting, but no comparable data have been published for Sub-Saharan Africa. When referenced to the Baka population itself, values for stunting were dramatically lower at 1.0% and 2.9% for 2-to-4 and 5-to-12 year olds, respectively. Wasting was also lower at 2.8% and 1.8% and was exceeded by obesity at 3.4% and 3.5%, respectively. Brachial perimeters and oedemas indicated rare severe malnutrition (< 2%) whilst moderate and severe anaemia were frequent (26.6% and 3.3%, respectively).

**Interpretation:** WHO child growth standards for stunting are clearly not applicable to Pygmies thus contradicting the widespread emphasis of their ethnicity-independent applicability. The inferred values for wasting and obesity are also difficult to interpret and are likely overestimated by the WHO criteria. To achieve UN Sustainable Development Goals and to fulfil our humanitarian responsibility for fellow man, we recommend that Pygmy specific growth standards are developed for the genetically differing Pygmy tribes.

## Introduction

Malnutrition refers to deficiencies, excesses or imbalances in a person’s intake of energy and/or nutrients. The term malnutrition covers ‘undernutrition’—which includes stunting (low height for age), wasting (low weight for height), underweight (low weight for age), micronutrient deficiencies or insufficiencies (a lack of important vitamins and minerals) and ‘overnutrition’ which includes overweight and obesity. Factors affecting undernutrition are manifold, ultimately determined by socio-economic and political factors that prolong poverty and social inequities[1,2]. The consequences of undernutrition include increased risk of infection, death, and delayed cognitive development resulting in low adult incomes and intergenerational transmission of poverty. Overnutrition is a major cause of noncommunicable diseases such as heart disease, stroke, diabetes and cancer.

Malnutrition is estimated to contribute to more than one third of all child deaths worldwide[3]. Despite a general reduction since the beginning of this century, numbers of undernourished children remain alarmingly high; in 2016, there were 155 million cases of stunting and 52 million cases of wasting in children under 5 years, corresponding to global rates of 23% and 8%, respectively[4]. Globally, almost two thirds of children suffering from malnutrition live in sub-Saharan Africa and Southern Asia. In sub-Saharan Africa, inadequate nutrition and poverty (almost half of the population in the region has an income of <$1.25 per day), which are interlinked, are widespread in rural and marginalised regions of the continent[5]. Assessments of child mortality in part caused by child malnutrition (including obesity) indicate that indigenous and tribal peoples are significantly disadvantaged compared to sympatric non-indigenous populations[6].

Diagnostic tools for assessing malnutrition are essential for paediatricians to target individual health, but also for local, national and international policy makers, politicians and development agencies to use as guidance instruments. More specifically, the identification of regional differences in the prevalence of local and national childhood malnutrition can help direct appropriate interventions in pursuance of the WHO global nutrition target of improving maternal, infant and young child nutrition by 2025[5].

The use of appropriate growth references is fundamental for reliably assessing malnutrition. In April 2006, the World Health Organization (WHO) released the WHO child growth standards, 16 years after a WHO working group on infant growth recommended that these standards should describe how children should grow rather than how they actually grow[7]. These standards, for children aged <5 years, are based on healthy, breastfed infants and children from healthy mothers without socio-economic and environmental constraints from six countries (Brazil, Ghana, India, Norway, Oman, USA). They are deemed good representatives for humankind, and are intended to be standards, not merely references, providing targets for children’s growth in all countries and for all ethnicities[8]. Whether this approach is appropriate for all people has been controversially discussed and a number of studies report national deviations from the WHO standards. For example, regardless of feeding practice, weights of Canadian children differed significantly for children with mothers of European ancestry compared with those with mothers of Asian/Pacific or South Asian heritage[9].

Among the standards, the indicator for stunting (length/height-for-age) is the most controversial[10]. Growth is under strong genetic regulation. The association between population genetic marker values and height has been confirmed in European populations, explaining about 25% of the observed variance[11]. In this socio-economically relatively homogeneous region, this means that three quarters of the variation in height is linked to non-genetic, environmental factors or genotype-by-environment interactions. However, the extent to which genetics determines height remains not fully understood as indicated by the much larger amount of about 80% of variance explained by genetic factors in twin studies[12]. The potential impact of socio-economic factors is also highlighted by upturns in adult height over the last century in certain countries e.g. an increase of 20.2 cm in South Korean women and 16.5 cm in Iranian men[10]. Moreover, a meta-analysis of over 11 million children aged <5 years from 55 countries and ethnic groups showed that study means for height were generally within ± 0.5 SD of the means in WHO growth standards, except for taller European populations and smaller populations in Saudi Arabia and Asian Indians[13]. Despite these clear differences, the WHO and others e.g. a recent editorial discussing the observed differences in Canadian children[9], claim that the WHO standards are adequate and “suitable for everyone”[14]. The justification for this claim is based on two premises. The first is that the population-specific deviation from the reference is relatively small compared to the cut-off points for stunting, wasting and underweight generally assessed using the z-score below minus two standard deviations (−2 SD) from the median of the WHO reference population[15]. The second is that any individual health assessment needs to be interpreted in the individual context and the debate of “which growth chart is less wrong”[14] appears academic. Accepting these arguments, a large number of countries have adopted the WHO standards with only a few, especially in Europe, not implementing them since 2011[8,15].

Whilst the WHO explicitly advises against country-or race-specific growth references[16], African Pygmies[17] (Note 1), which have a characteristically small body size (average adult height of 155 cm), merit explicit attention. Although the exact genetic mechanism remains tenuous, the Pygmy phenotype has genetic foundations impacting growth hormones and insulin-like growth factors[18]. Moreover, differences in growth patterns between Pygmy populations exist. For example, growth is faster during prenatal life in Eastern Pygmies such as the Efé and Sua, but slower during infancy in Western Pygmies, the Baka[18]. Baka growth rates are initially similar to non-Pygmies but then slow down significantly at about two years of age. However, adult Baka are similar in weight or even heavier than some Bantu populations in central Africa.

Pygmy-specific growth charts have not been developed, as suggested by some authors[13], nor have data been collected to evaluate the impact of their growth patterns on estimates of malnutrition. This information may help better understand the impact of ethnic background on malnutrition in countries such as Cameroon[6,19] where there is a high prevalence of stunting and wasting overall[5]. Life expectancy at birth for Baka Pygmies in Cameroon is an average of 35 years of age compared to the sympatric Bantu population who live about 22 years longer^6^. Such health differences between these two groups have been highlighted in the *Lancet* special issue on indigenous health[20].

In this study, we evaluate the applicability of WHO standards to quantify malnutrition in a sample of Pygmies populations living in south-eastern Cameroon. We assess stunting and wasting at an individual and population level. We also collected information on infection and anaemia rates to determine the overall health status of the subjects.

## Methods

### Data collection

Health data from Baka Pygmy children from the ages of one to 12 years old were collected in a total of 25 villages. Data were gathered in two separate campaigns in 2010 and 2018. We sampled 23 villages in 2010, and 10 villages in 2018. All villages are located within the Djoum-Mimtom road axis in the Dja et Lobo Department of southern Cameroon. The area is covered in tropical rainforests and borders Gabon and Congo.

Mobile clinics were set up in each village, under the auspices of Zerca y Lejos (ZyL), a Spanish NGO working on health and development issues in the region for over 15 years. ZyL has had a presence in all our study villages for all this time, where they regularly employ school teachers and support agricultural activities and health[21]. Our mobile clinics were open to all village inhabitants, who were encouraged to attend even if they were not unwell.

A total of 1073 persons were attended during the two campaigns. In 2010, we examined 550 (1-15 years old) Baka children, but in 2018, as many as 523 persons (from 6 months to >18 years) were assessed. During the latter campaign, we examined 404 Baka and 119 Bantu persons, out of which there were 200 children that were included in this study. Because we had population census data for the 10 villages sampled in 2018, we were able to calculate that between 30.3% and 90.3% (median 61.2%) of inhabitants were seen at the clinics.

A day before the clinic we held meetings with the village chiefs during which we obtained permission to perform out work. During the clinic, we verbally informed parents attending the clinics with their children of the purpose of the study and procedures used. Those who agreed to participate and to provide the data of their children for analyses gave written informed consent. Prior to data analysis, the names of participants were dissociated from all datasheets to maintain confidentiality. The Ministry of Science, Republic of Cameroon granted ethical approval.

For each child, we recorded data on village, ethnicity (self-declared; Baka and Bantu), weight (kg) and height (cm), clinical data on Hg (g l-1), brachial perimeter (mm) and absence/presence of oedema, and estimated age (in three-month intervals for babies up to 2 years old and in years for all others). Age estimation posed particular challenges since there was no formal certification of date of birth. Mothers were asked for the age of their children in years, since exact ages in months was not always possible. We crosschecked age estimations by dental eruption, but this method also only allows categorisation in years.

### Statistical analyses

We defined anaemia as a haemoglobin level two standard deviations below the mean for age according to WHO guidelines[22]. The web-based WHO Anthro Survey Analyser[23] was used to analyse anthropomorphic data of two to four year old Baka children with reference to the WHO Child Growth Standards. Henceforth we use H/A for height-for-age and W/L for weight-for-height. Observed data were compared to the growth potential as defined by WHO’s z-scores for healthy children. Cut-off criteria for z-scores for the identification of malnutrition in individuals were < −2 SD for wasting and stunting, < −3 SD for severe wasting and severe stunting, and > +2 SD for obesity.

We used the statistical software R[24] to further characterise the population-level measures for H/A and W/H in Baka children and to extend the analysis to five to 12 year old children, which is not covered by the WHO Anthro Survey Analyser[23]. We first age-and sex-corrected H/A and W/H values by subtracting the mean from all observations in each age and sex category retaining the same sign for uncorrected and corrected values. Skewness of the sample was statistically tested by a test for normality as implemented in the R package “normtest”[25] with 10,000 Monte Carlo Simulations.

## Results

### Anthropometrics

Height and weight measures were taken for a total of 685 children of 2 - 12 years; 2010 (*n*=460) and 2018 (*n*=213). H/A ratios changed rapidly from large values with large variances up to the age of about 7 years and then proceeded at more constant rate with lower variance (Figure 1). W/H ratios increased consistently with age with a slight increase in variances.

**Figure 1.**
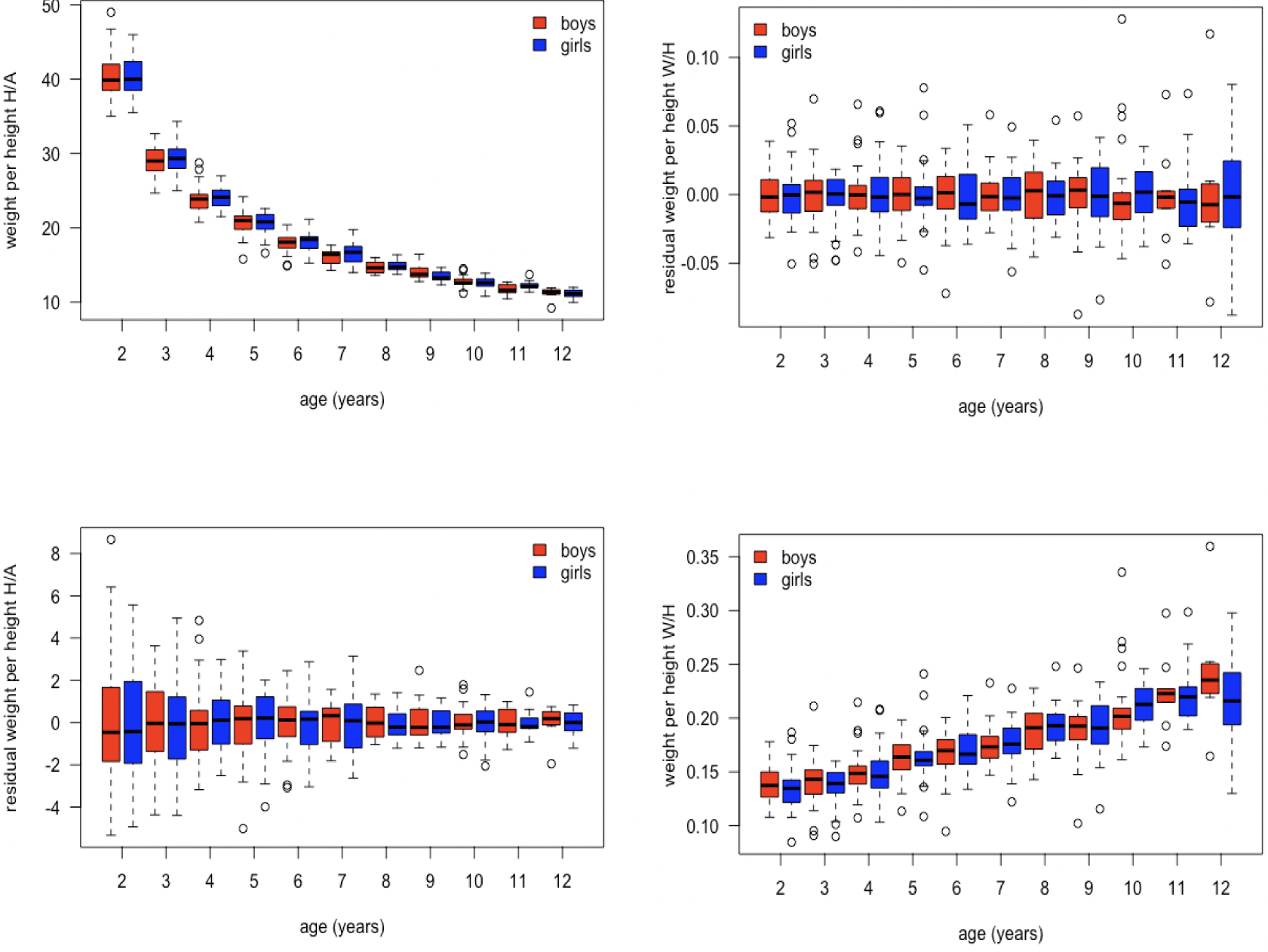
Height per age (top) and weight per height (bottom) in Baka Pygmy children aged two to twelve years.

Values for stunting, wasting and obesity were similar for boys and girls, and showed large overlaps of the 95% confidence intervals, calculated by the WHO Anthro Survey Analyzer (Table 1). Stunting, defined relative to the WHO child growth standard, was very high at 68.4% (95% CI: 62.8% - 73.5%) for 2- 4 year olds (Table 1). Almost every second child was severely stunted, 45.5% (39.8% - 51.3%). There was slightly more wasting at 8.2% (5.6% - 12.0%) than obesity at 6.5% (4.2%% - 10.0%). When using the distribution of H/A and W/H values of the study population to define malnutrition, values for stunting were dramatically lower than defined by the WHO standards for both age brackets with 1.0% for 2-4 year olds and 2.9% for 5-12 year olds. In each age class, the population-based obesity proportion was 2.4% and 3.9% higher than the respective level of wasting, which as 2.7% and 1.8%, respectively. Thus, population-based wasting was about a third of the value estimated with WHO standards.

**Table 1.** Summary of malnutrition in Baka children aged 2 to 12 years. The parameters wasting, stunting and obesity are defined relative to a reference population, which is the study population except for the WHO Anthro Survey Analyser, which applied the WHO child growth standard.

|  | 2-4 year olds | 2-4 year olds | 5-12 year olds |
| --- | --- | --- | --- |
| Reference population to define stunting, wasting & obesity | WHO growth standard | Population sample | Population sample |
| stunting |  |  |  |
| $\sum$ | 68.4 (62.8; 73.5), n=288 | 1.03, n=292 | 2.89, n=381 |
| ♀♀ | 62.9 (54.7; 70.5), n=143 | 1.40, n=143 | 2.20, n=182 |
| ♂♂ | 73.8 (66.0; 80.3), n=145 | 0.67, n=149 | 3.52, n=199 |
| wasting |  |  |  |
| $\sum$ | 8.2 (5.6; 12.0), n=291 | 2.75, n=291 | 1.84, n=380 |
| ♀♀ | 8.5 (4.8; 14.3), n=142 | 3.52, n=142 | 2.20, n=182 |
| ♂♂ | 8.1 (4.6; 13.7), n=149 | 2.01, n=149 | 1.52, n=198 |
| obesity |  |  |  |
| $\sum$ | 6.5 (4.2; 10.0), n=291 | 3.44, n=291 | 3.95, n=380 |
| ♀♀ | 4.9 (2.4; 10.0), n=142 | 3.52, n=142 | 4.40, n=182 |
| ♂♂ | 8.1 (4.6; 13.7), n=149 | 3.36, n=149 | 3.54, n=198 |

The distributions of measurements were not significantly different from normal distributions for girls in both age classes for H/A (Figure 2) and W/H (Figure 3), respectively. Significant skew was observed in boys for both H/A (Figure 2) and W/H (Figure 3) in both age classes. Despite these differences in skewness in boys and girls, the extent of the skew was in each case very small as indicated by the very small differences between means and medians (Figures 2 and 3). There was not only no bias towards stunting or wasting, but the population-based estimates for obesity exceeded wasting for boys in both age classes (Table 1).

**Figure 2.**
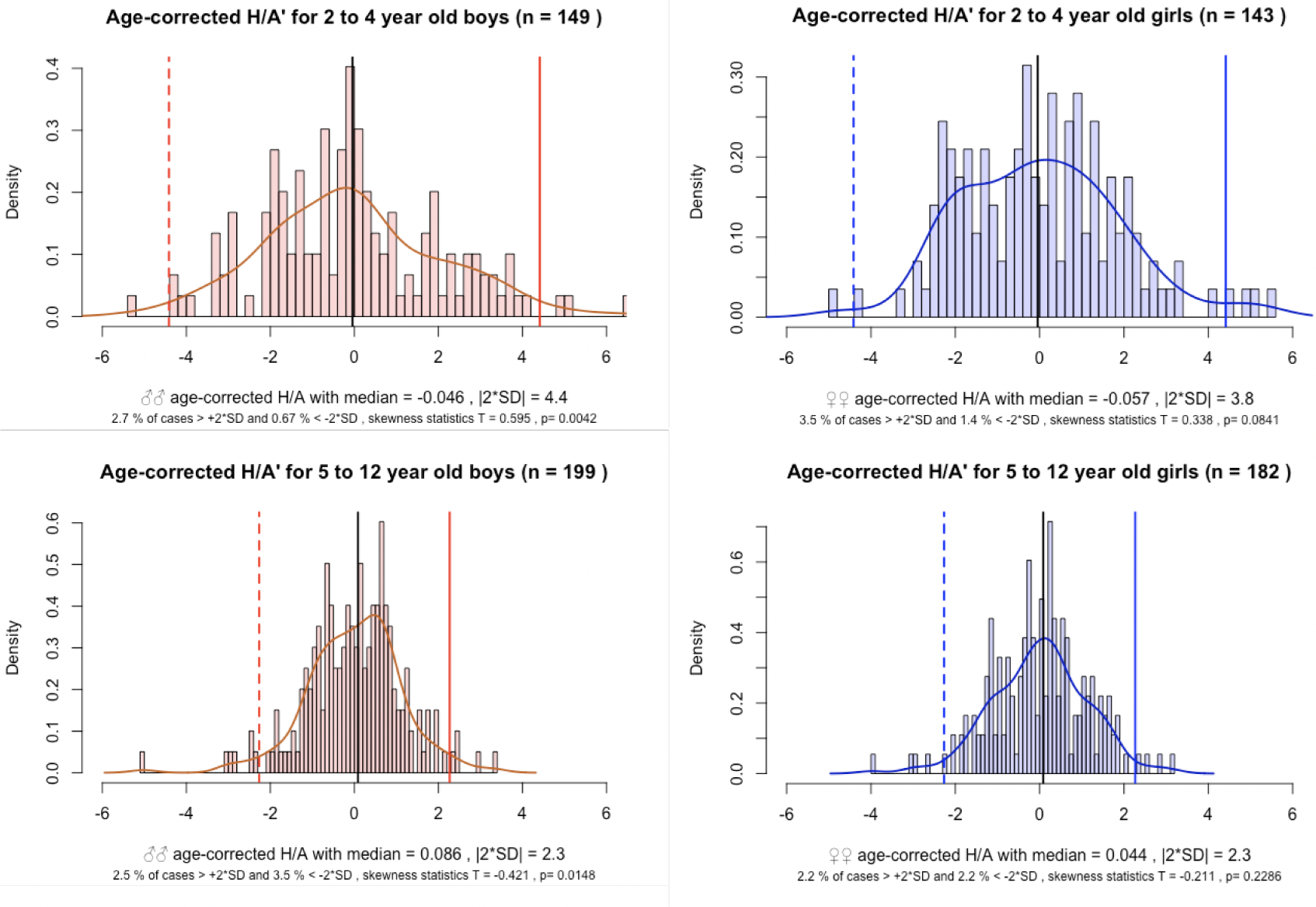
H/A scores and population-defined stunting Distributions, means, medians and ±2SD boundaries of age-corrected H/A scores for two-to-four year old (left) and five to-twelve year old right) Baka boys (top) and girls (bottom). Shown are the observed frequencies of age-corrected H/A scores (histograms), density distributions (solid curves), medians and ±2SD boundaries for each gender in each age-class. For each gender / age-class combination, percentages of cases where the ±2SD boundaries are exceeded, the test statistics T and resulting p-value of the skewness test are noted.

**Figure 3.**
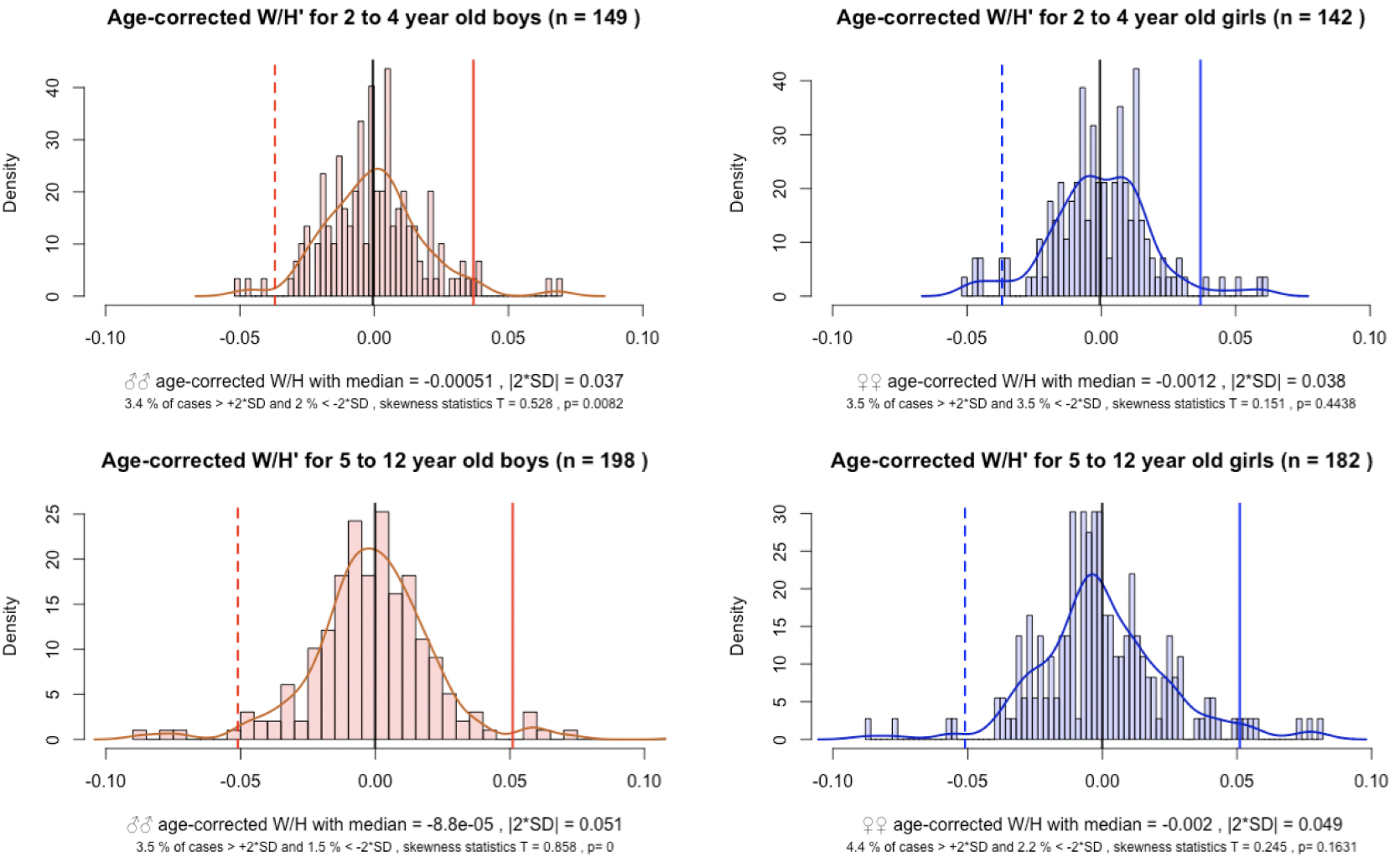
W/H scores and population-defined stunting. For details see description for Figure 2.

### Health assessments

Presence/absence of oedema was checked for 213 children; only one positive case (0.5%) was recorded. Brachial perimeters ranged from 116 mm to 195 mm (mean = 152.4 mm, median = 153 mm, *n* = 121) and only two cases (1.7%) were smaller than 125 mm. Haemoglobin concentrations indicated no anaemia for 293 children (46.7%, *n* = 628). Mild, moderate and severe anaemia was observed in 23.4%, 26.6% and 3.3% of children, respectively.

## Discussion

Sub-Saharan African countries remain exposed to the highest levels of child malnutrition[5]. In Cameroon, the average level of malnourishment ranks intermediate with 32.5% and 5.6% of reported stunting and wasting – defined based on WHO global standards - compared to averages of 28.8% (95% CI: 21.4 - 36.2) and 6.7% (95% CI: 4.2 - 9.2) for Central Africa[5]. Estimates of severe acute malnutrition in national surveys ranged from 1.9% in 2011 to 1.3% in 2014[5]. As in other sub-Saharan countries, Cameroon is ethnically highly diverse. However, how different ethnic groups differ with respect to malnutrition remains unknown. Studies have either concentrated on the dominant population segment, the ethnically diverse Bantu cluster[26], or have reported results from ethnically highly diverse health centres without including ethnicity in the analysis[27,28].

### Representativeness of the study population

The Baka Pygmies are a minority indigenous group that remain socio-economically highly disadvantaged. As a consequence, they suffer from considerably lower average health and shorter life expectancy compared to the majority sympatric Bantu population[6]. Since The Lancet’s recent focus on indigenous health, which spearheaded the predicament of the Baka[20], our study is the first to specifically present information on anthropometric indicators of Baka children. We are aware that our results may not be entirely representative of the whole Baka population because our data were taken from only one region where this group is currently found. Nonetheless, the sample coverage of our study villages was high, and our data represent the only information currently available on WHO growth standards for any Pygmy group. We confirm this on the basis of an extensive research we carried out of the peer-reviewed literature for Pygmies in general, prior to the study. Our study is also robust in that we focussed on general heath screening of children (and adults) and did not concentrate on malnourished individuals. The distribution of the data does not indicate any bias towards stunting in the population and there was even slightly more obesity than wasting observed, both supporting a reasonable representativeness of the study population to evaluate the adequacy of the WHO standards for Pygmies. The height data for 2 - 12 years indicate the obvious[18], namely that Baka children are on average small, but we challenge whether the WHO standards are representative and suitable to act as markers of their malnutrition.

### Meaningless stunting rates

A stunting rate, based on the WHO standards, of 68.4% for 2 - 4 year olds is astonishingly high compared to anywhere in the world (including Sub-Saharan Africa, where the rate was highest in Burundi in Eastern Africa with 57.7%)[5] and is, to the best of our knowledge, the highest globally recorded value. On the other hand, stunting when determined by the data from the population is only 1% for the same age class. The difference between these two estimates is extreme. What “real”, health-relevant levels of stunting are, remains, however, unclear. Height-per-age values are nearly normally distributed in Baka and do not indicate any skew towards relative shortness in the population (Figure 2). The exceptionally high WHO-based estimate is most likely caused by the by the growth curves of the Baka Pygmies, because their small phenotype has strong genetic foundations[18].

### Problematic wasting rates

Wasting in Baka 2 - 4 year olds was 8.2% higher than the 5.6% reported from the Cameroonian national survey, but lower than in 12 out of the 32 surveyed sub-Saharan African countries, where the highest level was reported for Niger with 18%[5]. However, similar to stunting, the results leave open the question of how representative is the anthropometric indicator for “wasting”. The percentage is possibly overestimated by the WHO child growth standards and underestimated by using the sample population itself as a reference. Even within Pygmies, there is variation in the genetic basis determining growth patterns as shown by the reported differences between Western and Eastern groups[18]. The impact on W/H has, however, not been determined. Like height-per-age, weight-per-height values are nearly normally distributed (Figure 3), producing similar percentages of wasting and obesity, respectively, for both, the application of the WHO standard and the Baka populations as reference. Shapes of distribution curves and obesity levels for Sub-Saharan African countries, have, however, not been considered in meta-analyses and are thus unavailable for comparison[5]. Considering all the additional indicators for malnutrition, especially the low life expectancy at birth for Baka in general and the high levels of anaemia in our study, the occurrence of obesity at similar levels as wasting is surprising and remains unexplained.

### Inadequacy of WHO growth standards for Pygmies

It is obvious that the specific case of Pygmies renders the often vehemently defended assumption that the WHO Child Growth Standards should be applied regardless of ethnicity[8,14–16] as inadequate for the height-per-age parameter. There is, of course, a logistic and economic imperative to apply the same child growth standards and to aim for the same model for optimal growth independently of ethnicity[15,16]. A number of studies have already argued that the WHO growth charts do not perfectly fit all combinations of ethnic/socio-economic background and ongoing secular trends, and no reference population or growth standard can provide an optimal statistical fit[14]. Thus, the problem of which approach to use seems to boil down to which method is ‘least wrong’. For the majority of populations under similar socio-economic situations, this question justifiably appears to matter little because the WHO growth standard seems to provide a reasonable standard[14]. However, the impact of the Pygmy phenotype on the identification of malnutrition, in particular stunting, is so extreme that the application of the WHO growth standard cannot be justified. Here, we also acknowledge that any ethnicity-based consideration for health care must be done with utmost concern for the highest ethical and moral standards, especially in ethnically diverse regions where ethnicity has played such a large role in human armed conflicts and persecutions. Possible or alleged genetic differences between human groups has been severely misused in the past and the consideration of such differences has provoked strong ethical concerns[29]. Although never expressed, these concerns might play a strong role in favouring a “seemingly inapplicable universal growth standard”. However, the WHO growth standards produce so vastly exaggerated stunting values and likely also bias other estimates of malnutrition that their application in Pygmy populations is not only obsolete but might be counterproductive.

### Consequences for development goals and individual health care

Given the uncertainty in interpreting the levels of observed wasting and obesity when applying WHO standards and or when using the within-population analysis, our data strongly indicate that the information gained from the analysis of anthropometric child data in Baka in particular and Pygmies in general is insufficient. It is therefore difficult to propose adequately targeted individual health care or clear national and international health policies for this ethnic group. Without a proper standard, the normally powerful parameter height-for-age will leave the clinician without the background information that is necessary to interpret these data[14].

On a population level, the current info on stunting and wasting in Baka are insufficient to provide meaningful information for achieving the UN Sustainable Development Goals for Baka, not to mention other Pygmy tribes which are even more data deficient. We also do not know how the growth patterns associated with the differing Pygmy genetic background affect national estimates of stunting and wasting not only in Cameroon but also in all the other West, Central and Eastern African countries with Pygmies, as genetic admixtures occur.

At an individual level, the application of the WHO standards in Pygmies opens the door for misinterpretation of the data by health practitioners and parents; the anthropometric variables can easily be wrongly understood as representing abnormal growth, and as a result mistaken or unnecessary solutions such as supplementary feeding prescribed. Considering that the central African region is under severe socio-economic pressures and focus of numerous politico-religious crises, armed conflicts and population displacements, health care is often overstretched making the necessary careful examination of anthropometric data less reliable. There are now concerted efforts to provide optimal health care to diminish the occurrence of severe acute malnutrition in Central Africa[28]. These laudable attempts could be disastrously undermined if misguided advice is given based on wrong growth standards.

The inadequacy of the UN growth standards for Pygmies highlights the urgent need to develop specific standards for ethnically and population genetically distinct groups, as already suggested in other studies[13]. If we are not to fail these more vulnerable and disadvantaged groups of people[5] [6,19] in areas that also likely to be affected by climate change[30], we need to act now to provide baseline data to evaluate children’s growth based on standards specific to their ethnic background.

## Contributors

JEF, BPG, NAP and GRB were responsible for the study design. SMB undertook the data analysis, data interpretation, figure construction, and manuscript writing. JEF and BPG contributed to data interpretation, and manuscript writing. AI contributed to the data interpretation, and manuscript writing. Data in the field were gathered by BPG, NAP, MAA, YHS, RP, GRB, MA, EAM, RO, BAZ, AMC, CGS, CRLG, FLRS, HA, and IAR. All authors approved the final version to be published.

## Acknowledgments

We thank Maria Rebollo for allowing us to use data gathered during the 2010 campaign. Julienne Meyina and Mirabelle Assampelle, nurses within the Zerca y Lejos health programme, assisted during the campaigns. Funding was provided by the UK Darwin Initiative (Project 24029).

## Note 1. Use of the term Pygmy

Although numerous alternative terms to Pygmy have been used to refer the rainforest hunter-gatherers of the Congo Basin, none have been agreed upon by academics or the people themselves to replace it. Although some academics and Central African government officers feel the term Pygmy is derogatory or does not adequately represent the people, the term Pygmy *sensu lato*, to refer to all hunter-gatherer groups in Central Africa, is widely used by a broad group of people in Europe, Japan, the United States and Africa. Moreover, International and local NGOs use the term in their titles or literature e.g. Pygmy Survival Alliance, Forest Peoples’ Programme. Survival International, Rainforest Foundation, Reseau Recherches Actions Concerteees Pygmees, Centre d’Accompagnement des Autochtones Pygmees et Minoritaires Vulnerables and the Association for the Development of Pygmy Peoples of Gabon. Congo Basin conservation groups, such as World Wildlife Fund and Wildlife Conservation Society and international human rights groups working in the region, such as UNICEF and Integrated Regional Informaton Networks (IRIN), also regularly use the term Pygmy in their literature.

We consider all groups under the umbrella of Pygmy as expressing an equivalent spatial relationship between their presence and their immediate environment. In so doing, we do not ignore the fact that various ‘Pygmy’ groups express distinct cultures and in some case ethnicity from other ‘Pygmy’ groups, and is not meant in any way a disrespect to the various ethnicities. Although it is likely that there may be cultural reasons for geographical location and distribution, we argue that the ecological setting is a primary driver in humans in choosing localities to live in. Ichikawa (2014)14:332-3 – ‘The forest plants and animals provide the people with the basis for their cultural identity. Their life and culture cannot be maintained without the forest in its entirety … The destruction of the forest would result in the deterioration of a culture that is heavily dependent on, and in very significant ways a part of, the Congo Basin rainforest.’

## References

1. Heaton TB, Crookston B, Pierce H, Amoateng AY. Social inequality and children’s health in Africa: a cross sectional study. International Journal for Equity in Health. 2016;15. doi:10.1186/s12939-016-0372-2

2. Pridmore P, Carr-Hill R. Tackling the drivers of child undernutrition in developing countries: what works and how should interventions be designed? Public Health Nutrition. 2011;14: 688–693. doi:10.1017/S1368980010001795

3. WHO. Global Database on Child Growth and Malnutrition. In: World Health Organization [Internet]. 2018 [cited 5 Jul 2018]. Available: http://www.who.int/nutgrowthdb/about/en/

4. United Nations. The Sustainable Development Goals Report 2017. New York: United Nations; 2017.

5. Akombi BJ, Agho KE, Merom D, Renzaho AM, Hall JJ. Child malnutrition in sub-Saharan Africa: A meta-analysis of demographic and health surveys (2006-2016). Wieringa F, editor. PLOS ONE. 2017;12: e0177338. doi:10.1371/journal.pone.0177338

6. Anderson I, Robson B, Connolly M, Al-Yaman F, Bjertness E, King A, et al. Indigenous and tribal peoples’ health (The Lancet–Lowitja Institute Global Collaboration): a population study. The Lancet. 2016;388: 131– 157. doi:10.1016/S0140-6736(16)00345-7

7. Bloem M. The 2006 WHO child growth standards. 2007;334: 705–706. doi:10.1136/bmj.39155.658843.BE

8. WHO Multicentre Growth Reference Study Group. WHO Child Growth Standards based on length/height, weight and age: WHO Child Growth Standards. Acta Paediatrica. 2006; Suppl 145: 76–85. doi:10.1111/j.1651-2227.2006.tb02378.x

9. Park AL, Tu K, Ray JG, the Canadian Curves Consortium. Differences in growth of Canadian children compared to the WHO 2006 Child Growth Standards. Paediatric and Perinatal Epidemiology. 2017;31: 452–462. doi:10.1111/ppe.12377

10. NCD Risk Factor Collaboration (NCD-RisC). A century of trends in adult human height. eLife. 2016;5. doi:10.7554/eLife.13410

11. Robinson MR, Hemani G, Medina-Gomez C, Mezzavilla M, Esko T, Shakhbazov K, et al. Population genetic differentiation of height and body mass index across Europe. Nature Genetics. 2015;47: 1357–1362. doi:10.1038/ng.3401

12. Jelenkovic A, Sund R, Hur Y-M, Yokoyama Y, Hjelmborg J v. B, Möller S, et al. Genetic and environmental influences on height from infancy to early adulthood: An individual-based pooled analysis of 45 twin cohorts. Scientific Reports. 2016;6. doi:10.1038/srep28496

13. Natale V, Rajagopalan A. Worldwide variation in human growth and the World Health Organization growth standards: a systematic review. BMJ Open. 2014;4: e003735. doi:10.1136/bmjopen-2013-003735

14. Ong KK. WHO Growth Standards - Suitable for Everyone? Yes. Paediatric and Perinatal Epidemiology. 2017;31: 463–464. doi:10.1111/ppe.12396

15. de Onis M, Onyango A, Borghi E, Siyam A, Blössner M, Lutter C. Worldwide implementation of the WHO Child Growth Standards. Public Health Nutrition. 2012;15: 1603–1610. doi:10.1017/S136898001200105X

16. de Onis M. Child Growth and Development. In: Semba RD, Bloem MW, editors. Nutrition and health in developing countries. 2nd ed. Totowa, NJ: Humana Press; 2008. pp. 133–138.

17. Olivero J, Fa JE, Farfán MA, Lewis J, Hewlett B, Breuer T, et al. Distribution and Numbers of Pygmies in Central African Forests. Caramelli D, editor. PLOS ONE. 2016;11: e0144499. doi:10.1371/journal.pone.0144499

18. Ramírez Rozzi FV, Koudou Y, Froment A, Le Bouc Y, Botton J. Growth pattern from birth to adulthood in African pygmies of known age. Nature Communications. 2015;6. doi:10.1038/ncomms8672

19. Ohenjo N, Willis R, Jackson D, Nettleton C, Good K, Mugarura B. Health of Indigenous people in Africa. The Lancet. 2006;367: 1937–1946. doi:10.1016/S0140-6736(06)68849-1

20. The Lancet. Indigenous health: a worldwide focus. The Lancet. 2016;388: 104. doi:10.1016/S0140-6736(16)31020-0

21. Zerca y Lejos [Internet]. [cited 24 Oct 2018]. Available: http://zercaylejos.org/?lang=en

22. World Health Organization. Haemoglobin concentrations for the diagnosis of anaemia and assessment of severity. 2011;

23. WHO. The WHO Anthro Survey Analyser. In: World Health Organization [Internet]. 2018 [cited 5 Jul 2018]. Available: https://whonutrition.shinyapps.io/anthro/

24. R Foundation for Statistical Computing. R [Internet]. 2016. Available: https://www.r-project.org

25. Gavrilov I, Pusev R. Package ‘normtest.’ 2015.

26. Nagahori C, Kinjo Y, Tchuani JP, Yamauchi T. Malnutrition among vaccinated children aged 0-5 years in Batouri, Republic of Cameroon. Journal of General and Family Medicine. 2017;18: 365–371. doi:10.1002/jgf2.104

27. Chiabi A, Malangue B, Nguefack S, Dongmo FN, Fru F, Takou V, et al. The clinical spectrum of severe acute malnutrition in children in Cameroon: a hospital-based study in Yaounde, Cameroon. Translational Pediatrics. 2017;5: 32–39. doi:10.21037/tp.2016.07.05

28. Ndzo JA, Jackson A. Outcomes of children aged 6–59 months with severe acute malnutrition at the GADO Outpatient Therapeutic Center in Cameroon. BMC Research Notes. 2018;11. doi:10.1186/s13104-018-3177-0

29. Winegard B, Winegard B, Boutwell B. Human Biological and Psychological Diversity. Evolutionary Psychological Science. 2017;3: 159–180. doi:10.1007/s40806-016-0081-5

30. Berrang-Ford L, Dingle K, Ford JD, Lee C, Lwasa S, Namanya DB, et al. Vulnerability of indigenous health to climate change: A case study of Uganda’s Batwa Pygmies. Social Science & Medicine. 2012;75: 1067– 1077. doi:10.1016/j.socscimed.2012.04.016

